# Extensive Transcriptional and Translational Regulation Occur During the Maturation of Malaria Parasite Sporozoites

**DOI:** 10.1101/642298

**Authors:** Scott E. Lindner, Kristian E. Swearingen, Melanie J. Shears, Michael P. Walker, Erin N. Vrana, Kevin J. Hart, Allen M. Minns, Photini Sinnis, Robert L. Moritz, Stefan H.I. Kappe

**Affiliations:** Department of Biochemistry and Molecular Biology, The Huck Center for Malaria Research, Pennsylvania State University, University Park, PA; Institute for Systems Biology, Seattle, WA; Department of Molecular Microbiology & Immunology, Johns Hopkins Bloomberg School of Public Health, Baltimore, MD; Center for Global Infectious Disease Research, Seattle Children’s Research Institute, Seattle, WA; Department of Global Health, University of Washington, Seattle, WA

**Keywords:** Plasmodium, sporozoite, translational repression, RNA-seq, proteomics

## Abstract

*Plasmodium* sporozoites are transmitted from an infected mosquito to mammals in which they infect the liver. The infectivity profile of sporozoites changes as they egress from oocysts on the mosquito midgut into the hemocoel, and then invade the salivary glands, where they maintain a poised and infectious state until transmission occurs. Upon transmission, the sporozoite must then navigate the host skin, vasculature, and liver. All of these feats require distinct repertoires of proteins and capabilities that are coordinated in an appropriate temporal manner. Here, we report the comprehensive and dynamic transcriptomes and proteomes of both oocyst sporozoite and salivary gland sporozoite stages in both rodent-infectious *Plasmodium yoelii* parasites and human-infectious *Plasmodium falciparum* parasites. These data robustly define mRNAs and proteins that are <u>U</u>pregulated in <u>O</u>ocyst <u>S</u>porozoites (UOS) or <u>U</u>pregulated in Infectious <u>S</u>porozoites (UIS), which include critical gene products for sporozoite functions, as well as many of unknown importance that are similarly regulated. Moreover, we found that *Plasmodium* uses two overlapping, extensive, and independent programs of translational repression across sporozoite maturation to temporally regulate specific genes necessary to successfully navigate the mosquito vector and mammalian host environments. Finally, gene-specific validation experiments of selected, translationally repressed transcripts in *P. yoelii* confirmed the interpretations of the global transcriptomic and proteomic datasets. Together, these data indicate that two waves of translational repression are implemented and relieved at different times in sporozoite maturation to promote its successful life cycle progression.

## Introduction

Malaria remains one of the great global health problems today, taking a large toll on people in the tropics and subtropics. This disease, caused by *Plasmodium* parasites, affects over 200 million people annually and kills over 400,000 (WHO World Malaria Report 2018). While a protein-based subunit vaccine (RTS,S) has recently been licensed and is being used for pilot implementation in three Sub-Saharan African countries, its protection has been limited and relatively short-lived in clinical trials^1^. Identifying and fully testing an effective and long-lasting malaria vaccine remains a chief goal that has yet to be achieved. Accomplishing this will require greater knowledge of the basic biology of the pre-erythrocytic sporozoite and liver stage parasites. Promising new whole-parasite vaccine candidates, based upon the sporozoite form of the parasite, are on the horizon and hopefully will meet these needs^2^.

*Plasmodium* parasites are transmitted between hosts by an infected female *Anopheles* mosquito (reviewed in^3^). Following uptake of male and female gametocytes by the mosquito during a blood meal of an infected host, these parasites activate into male and female gametes in the midgut, fuse into a zygote, and develop into a motile ookinete, which burrows through the midgut wall and establishes an oocyst under the basal lamina. Within the oocyst, the parasite undergoes sporogony to produce up to five thousand oocyst sporozoites^4^. These oocyst sporozoites are weakly infectious if injected directly into a naïve mammalian host^5^, but become highly infectious following proteolytic rupture of the oocyst wall and transit through the mosquito hemocoel. Sporozoites further gain infectivity after invasion of the salivary glands^5, 6^. Interestingly, the gain of infectivity for the mammalian host is concomitant with a loss of infectivity for the salivary glands. Within the glands, the sporozoites await transmission as salivary gland sporozoites, which occurs when the mosquito takes its next blood meal. Infection of a host begins with the injection of salivary gland sporozoites into the skin. Following this, sporozoites exit the bite site in the skin, locate and enter the vasculature, and passively travel to the liver, where sporozoites can productively infect hepatocytes, which initiates the life cycle progression in the mammalian host^7^. Moreover, because relatively few sporozoites are injected during a mosquito bite^8^, this transmission event bottleneck has been the focus of intervention efforts using drugs, subunit vaccines, and attenuated whole parasite vaccines^2^.

Fundamental studies of sporozoite biology have informed efforts to inhibit and/or arrest the parasite during development. To date, genetically-attenuated parasite (GAP) vaccine candidates were generated by the deletion of genes whose transcripts are Up-regulated in Infective (Salivary Gland) Sporozoites (UIS). The UIS gene family was originally determined by a subtractive cDNA hybridization approach^9^, where 30 expressed sequence tags (ESTs) were identified as being more abundant in salivary gland sporozoites compared to oocyst sporozoites. Of these, 23 map to currently annotated genes, with several of the original ESTs mapping to the top hit (UIS1). With the advent of microarray-based transcriptomics, a renewed effort to identify both UIS and Up-regulated in Oocyst Sporozoites (UOS) genes that may be similarly exploited to produce effective GAP vaccines identified 124 UIS and 47 UOS genes^10^. Interestingly, only 7 of the original 23 UIS genes were confirmed in this expanded study. However, these UIS genes (UIS1, UIS2, UIS3, UIS4, UIS7, UIS16, UIS28) have proven to encode some of the most important proteins for the transmission and transformation of the sporozoite into a liver stage parasite, as well as for liver stage development^11–14^.

In addition to transcriptional controls placed upon key transcripts, the malaria parasite also imposes translational repression upon specific mRNAs in female gametocytes, and for at least a few mRNAs in salivary gland sporozoites (reviewed in^15, 16^). This model allows for the proactive production of mRNAs and restriction of their translation before transmission, and yet enables a strategy of just-in-time production of these proteins after transmission when they are needed. However, this strategy (high transcription, low/no translation) is energetically costly, and model eukaryotes and human cells have evolved to avoid this gene regulatory combination (“Crick Space”) except in specific, beneficial situations^17^. In light of this, it is notable that *Plasmodium* parasites have evolved to use translational repression for transmission to the mosquito, which has been clearly seen for mRNAs (*e.g. p28*) in female gametocytes^18^. The mechanisms underlying this have been established most thoroughly in *P. berghei*, where DOZI (a DEAD-box RNA helicase orthologous to human DDX6) and CITH (an Lsm14 orthologue) bind, stabilize, and translationally repress specific mRNAs in female gametocytes^18–20^. Recently, the extent of translational repression in *P. falciparum* female gametocytes was assessed by mass spectrometry-based proteomics and RNA-seq^21^. In this stage, the parasite expresses over 500 transcripts with no evidence for their encoded proteins, with over half of these maternal gene products being uncharacterized. Enriched in this set of translationally repressed mRNAs are those that encode for functions needed post-transmission and include gene families previously shown to be translationally repressed. These data support the model that large-scale preparations are made by female gametocytes, despite high energetic costs, to prepare and await transmission by storing and protecting specific mRNAs needed to establish the infection of the mosquito. However, the extent and precise mechanisms of how translational repression is imposed in sporozoites has not been established beyond targeted studies that suggest that the PUF2 RNA-binding protein may act upon *cis* elements found within the coding sequence of the *uis4* mRNA^13^.

Many previous efforts that aimed to determine the global transcriptomes and proteomes of *Plasmodium* sporozoites were greatly restricted due to substantial contamination with material from the mosquito vector and its microbiome in these samples^22–24^. In response to this, we have developed a scalable, discontinuous density gradient purification approach for sporozoites that greatly reduces contamination from the mosquito and its microbes^25^. The resulting fully infectious sporozoites have allowed extensive ChIP, transcriptomic (RNA-seq) and proteomic (nanoLC-MS/MS) analyses of sporozoites from both rodent-infectious (*P. yoelii*) and human-infectious (*P. falciparum*, *P. vivax*) parasites^26–32^. Taken together, these studies demonstrate that ‘omics’ level efforts on sporozoites are now experimentally practical, and thus reopen long standing questions of mechanisms underlying critical sporozoite functions.

Here we have addressed one of the foremost of these questions: how and when does the sporozoite prepare molecularly for transmission from the mosquito vector to the mammalian host? Using RNA-seq-based transcriptomics and nanoLC-MS/MS-based proteomics, we have characterized both oocyst sporozoites and salivary gland sporozoites of both rodent-infectious (*P. yoelii*) and human-infectious (*P. falciparum)* species. Together, these data provide a comprehensive assessment of mRNA and protein abundances, provide evidence for extensive post-transcriptional regulation of the most abundant mRNAs, and demonstrate that two distinct and likely orthogonal translational repression programs are active during sporozoite maturation.

## Results

### Dynamic Transcriptional Regulation during Plasmodium Sporozoite Development

Important insights into how sporozoites mature and become infectious were gained from studies of the Upregulated in Oocyst Sporozoites (UOS) and Upregulated in Infectious Sporozoites (UIS) transcripts in *Plasmodium*. Moreover, a number of these UIS genes turned out to be essential to hepatocyte infection and the early liver stage parasite. However, prior studies were limited by the methods and instrumentation available, thus resulting in an incomplete view of transcriptional regulation in the sporozoite. By leveraging RNA-seq and greatly improved sporozoite purification strategies, we could now achieve a more comprehensive transcriptome and differential expression analyses of sporozoites from rodent-infectious (*P. yoelii*) and human-infectious (*P. falciparum*) species. Additionally, as the sporozoite undergoes a transition from being weakly infectious to highly infectious for the mammalian host, which occurs while in transit through the hemocoel from the oocyst to the salivary glands and within the salivary glands^5, 6^, we assessed both the oocyst sporozoite and salivary gland sporozoite transcriptomes (Figure 1, Table 1, Table S1).

**Figure 1:**
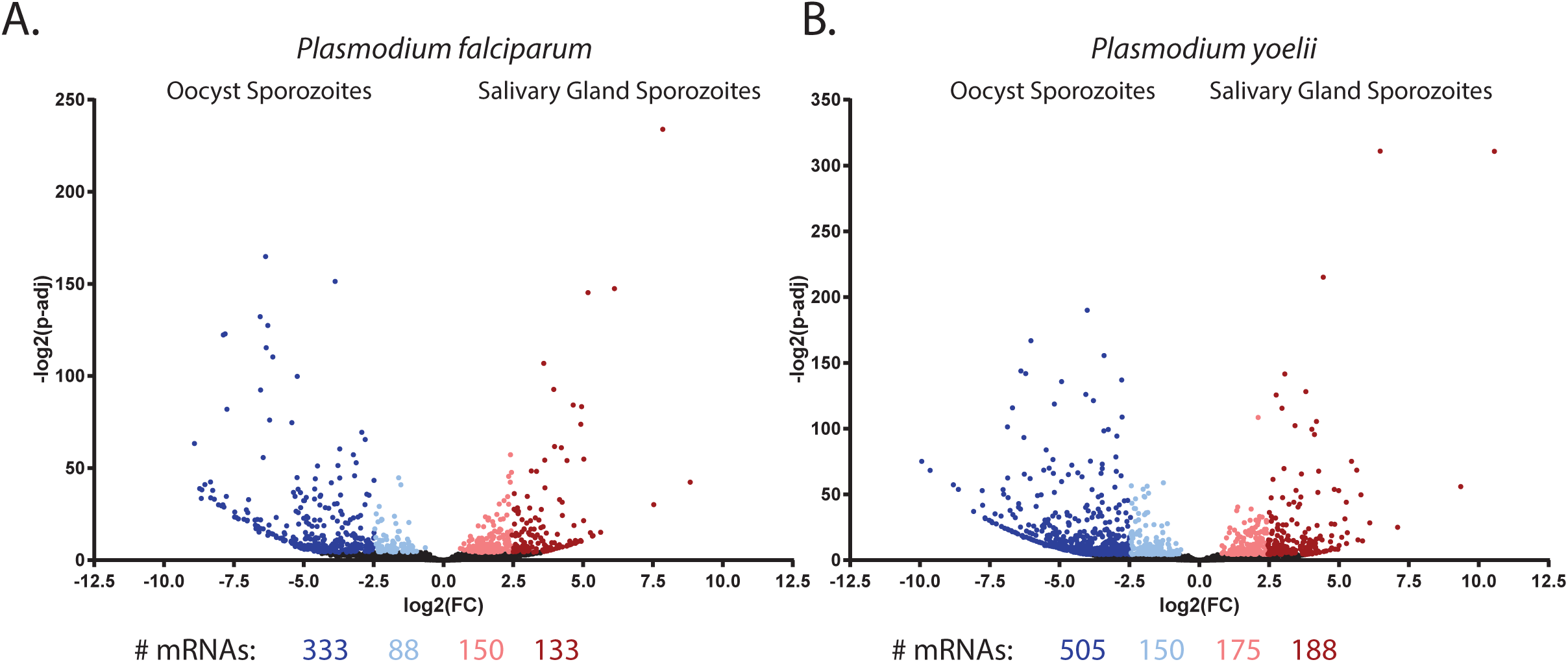
Comparative transcriptomics of oocyst and salivary gland sporozoites. RNA from purified (A) *P. falciparum* or (B) *P. yoelii* sporozoites isolated from oocysts or the salivary gland was assessed by RNA-seq, and transcript abundances compared by DEseq2. Transcripts were binned based upon fold change and adjusted p-value, with mRNAs shaded in lighter (+/-1 to 2.5 log2 fold change) or darker shades (>2.5 log2 fold change).

**Table 1:**
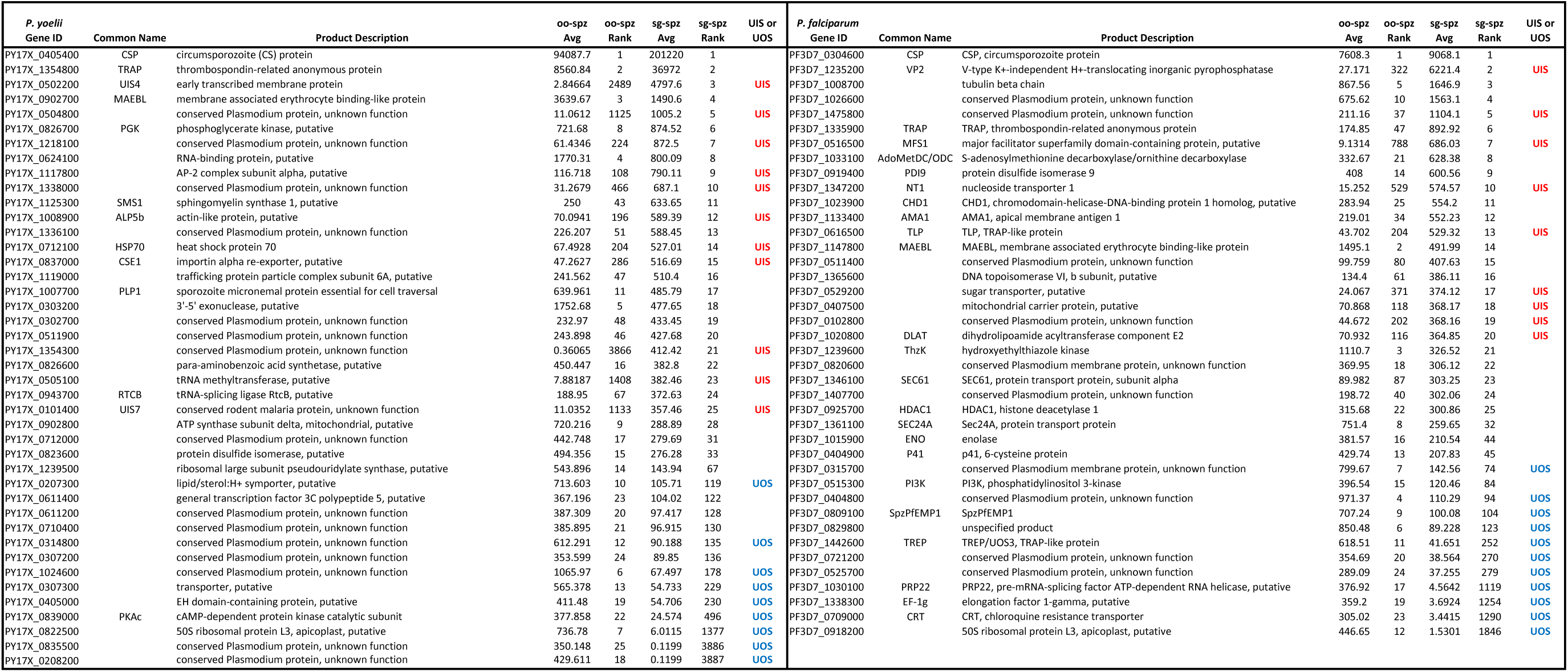
The top 25 most abundant mRNAs in oocyst sporozoites and salivary gland sporozoites of *P. falciparum* and *P. yoelii*. Transcripts are defined as Upregulated in Oocyst Sporozoites (UOS) or Upregulated in Infectious Sporozoites (UIS) mRNAs as well when their abundance is in the top decile, and their fold change is >6-fold in one sporozoite stage compared to the other. A complete listing of mRNA abundances and fold changes between sporozoite stages is provided in Table S1.

First, using *P. yoelii* (17XNL non-lethal strain) rodent-infectious parasites we identified 4195 and 3887 RNAs with detectable and unambiguous sequence reads present in *P. yoelii* oocyst sporozoites and salivary gland sporozoites, respectively. Similarly, with *P. falciparum* (NF54 strain) human-infectious parasites, we identified 3535 and 3575 detectable and unambiguous RNAs in oocyst sporozoite and salivary gland sporozoite stages, respectively. Many well-characterized genes were among the most abundant transcripts in these two stages, including *apical membrane antigen 1 (ama1), circumsporozoite protein (csp)*, *membrane-associated erythrocyte binding-like protein* (*maebl)*, *perforin-like protein 1 (plp1/pplp1/spect2)*, *thrombospondin-related anonymous protein (trap)*, *trap-like protein (tlp), upregulated in infectious sporozoites 4 (uis4)*, and others (Table 1, Table S1)^33–38^. However, numerous mRNAs encode for proteins with un/under-characterized roles in the parasite. Notably, several of the most abundant mRNAs in oocyst and salivary gland sporozoites in both species encode for uncharacterized proteins, some of which (e.g. *py17x_0208200*, *py17x_0835500*, *py17x_1354300*) undergo the same extreme swings in transcript abundance between these stages as does *pyuis4* (>1000-fold). Finally, we found that a recently described sporozoite var gene (SpzPfEMP1) is robustly expressed in not only oocyst sporozoites as previously reported, but also in salivary gland sporozoites and thus may simplify the recently described model of how this interesting var gene is regulated (Table 1)^32^. Given the transcript abundance of the novel and uncharacterized genes of these lists, they warrant a prioritized assessment.

Previous definitions of UOS or UIS mRNAs were assigned using lower thresholds of >2-fold increases in transcript abundance for any detectable transcript between oocyst and salivary gland sporozoites, which in part were dictated by the power of subtractive cDNA hybridization or microarray approaches available at the time^9, 10^. With greatly improved approaches, we have made the definitions of UIS and UOS RNAs more stringent by assigning thresholds whereby transcripts must be both in the top decile of abundance, and must be >6-fold more abundant in one stage compared to the other. Using these parameters, we have defined 164 UOS mRNAs and 88 UIS mRNAs in *P. yoelii*, and 101 UOS mRNAs and 68 UIS mRNAs in *P. falciparum* (Table S2). Few of the UOS transcripts previously defined remain so using these more stringent thresholds, but robustly include the previous top UOS hit: TREP/UOS3^10^.

Additional UOS transcripts include those that encode for proteins important for sporozoite functions in the mosquito and the initial infection of the new host, and include those that encode for RNA metabolic processes, protein translation, heat shock proteins (HSP20), the glideosome/inner membrane complex (GAPM3, IMC1m), vesicular trafficking, and transporters. Similarly, a core set of the most abundant UIS transcripts remain defined as such here: *uis1*, *uis2*, *uis3*, *uis4*, *uis7*, *uis8*, and *uis28*. Strikingly, *pyuis4* transcript abundance increases 1500-fold and reaffirms the use of its promoter for highly enriched expression of transgenes in salivary gland sporozoites^28, 39^. An additional 71 *P. yoelii* and 53 *P. falciparum* transcripts that were not in the top decile of RNA abundance, but were in the seventh to ninth decile, increase >10-fold in abundance in salivary gland sporozoites vs. oocyst sporozoites. These include *pfccr4*, the *pfdbp10* and *pydbp10* RNA helicase, *pfslarp/pfsap1* (40-fold), *pynek1* (38-fold), *pypplp3*, *pyplp5*, *pfptex150*, *pyuis5* (22-fold) and *pyuis12* (40-fold) (Table S3). Beyond the previously defined UIS mRNAs, several other transcripts are notable as their protein products are or may be important to the function of salivary gland sporozoites: FAS-II pathway proteins, fatty acid modifiers, plasma membrane transporters, adhesins/surface proteins (*p113*, *speld*, *tlp*), traversal-related proteins (*celtos*), heat shock proteins, and ApiAP2 specific transcription factors (*e.g. py17x_0523100, pf3d7_0420300*)^40–47^. Together, these transcripts encode for proteins that encompass many essential attributes necessary for sporozoite development, transmission, and infectivity. However, it is striking that while a similar regulatory strategy is employed by sporozoites of both parasite species, there is less overlap in the transcripts that are regulated than might be expected, especially in oocyst sporozoites. These findings underscore the strengths and importance of using comparative approaches with human- and rodent-infectious species to identify the important and conserved molecular components of infection.

### Proteomic Comparisons of Plasmodium Sporozoites

While transcriptomics can provide an important window into gene expression, the inclusion of proteomics provides a much more comprehensive understanding of the parasite and its functions. To determine the presence and steady-state abundance of proteins found within *Plasmodium* sporozoites, the global proteomes of both *P. yoelii* and *P. falciparum* oocyst sporozoites were determined by nanoLC/MS/MS and were compared to our previously published salivary gland sporozoite global proteomes^29^. This approach (steady-state protein abundance) was used, as it is compatible with currently feasible sporozoite production and purification capabilities, whereas ribosome profiling remains technically unfeasible with sporozoite samples due to the number of highly purified sporozoites that are required. Together, these four datasets now allow a more complete understanding of the oocyst and salivary gland sporozoite stages, and also allows for the definition of UOS Proteins and UIS Proteins for these two species of the malaria parasite.

Using approximately four million purified sporozoites per biological replicate, protein lysates were separated in a single lane of a gradient SDS-polyacrylamide gel, digested with trypsin, and the resulting tryptic peptides were extracted and subjected to nanoLC-MS/MS. Resulting mass spectra were assessed with the Trans-Proteomic Pipeline (TPP) to identify peptides and to infer identities of proteins. In sum, re-analysis of our previously acquired *P. falciparum* data identified 2037 salivary gland sporozoite proteins^29^ and we now here also identify 1430 oocyst sporozoite proteins; similarly, from our previously acquired *P. yoelii* data, we identified 1773 salivary gland sporozoite proteins, and here identify 1760 oocyst sporozoite proteins (Figure 2, Table 2, see Table S1 for a complete list). In addition to detecting well-characterized sporozoite proteins (CSP, CelTOS, TRAP, IMC/glideosome proteins, ALBA proteins, SIAP1) and abundant housekeeping proteins (histones, HSPs, GAPDH, translation-related proteins), dynamic changes in the abundance of specific proteins between these sporozoite stages were also identified. For instance, in both *P. yoelii* and *P. falciparum*, CelTOS, GEST and SPELD were not detected or were only weakly expressed in oocyst sporozoites, but were among the most abundant proteins in salivary gland sporozoites (Table 2, Table S1). Moreover, the presence/absence of cellular regulators such as specific ApiAP2s, histone modifiers, RNA-binding proteins and other proteins (Table S1) agree with previous reports describing how these types of regulation may be used by the sporozoite^32, 40, 48^.

**Figure 2:**
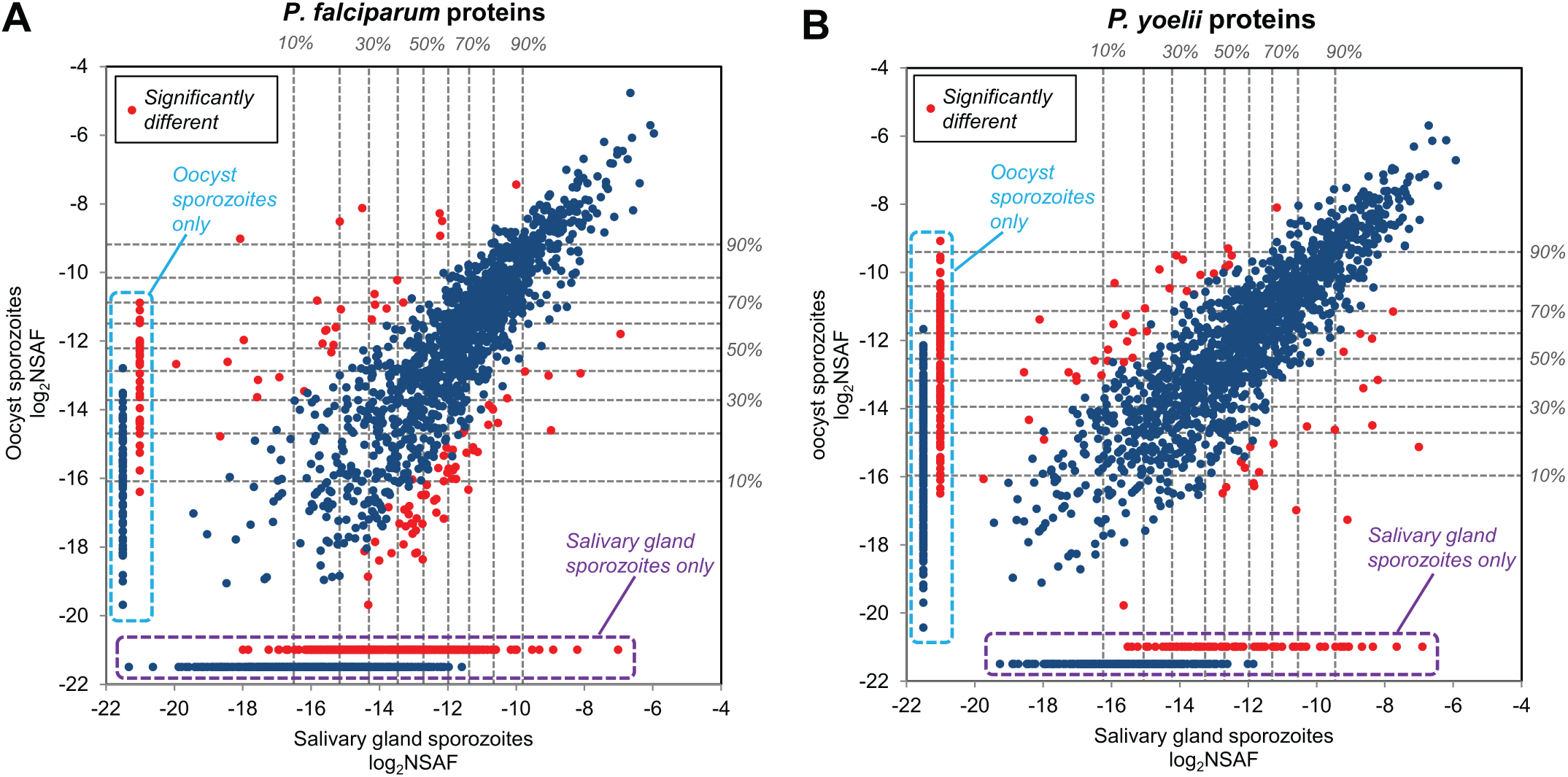
Comparative total proteomics of oocyst sporozoites and salivary gland sporozoites. Total protein from purified (A) *P. falciparum* or (B) *P. yoelii* sporozoites isolated from oocysts or the salivary gland was separated by a gradient SDS-PAGE, digested with trypsin, extracted, and subjected to analysis. Proteins with statistically significant differences in abundance are noted with red dots, and expression percentile thresholds are provided by dashed lines. Proteins that were only detected in oocyst sporozoites or salivary gland sporozoites are boxed in light blue and purple dashed lines, respectively.

**Table 2:**
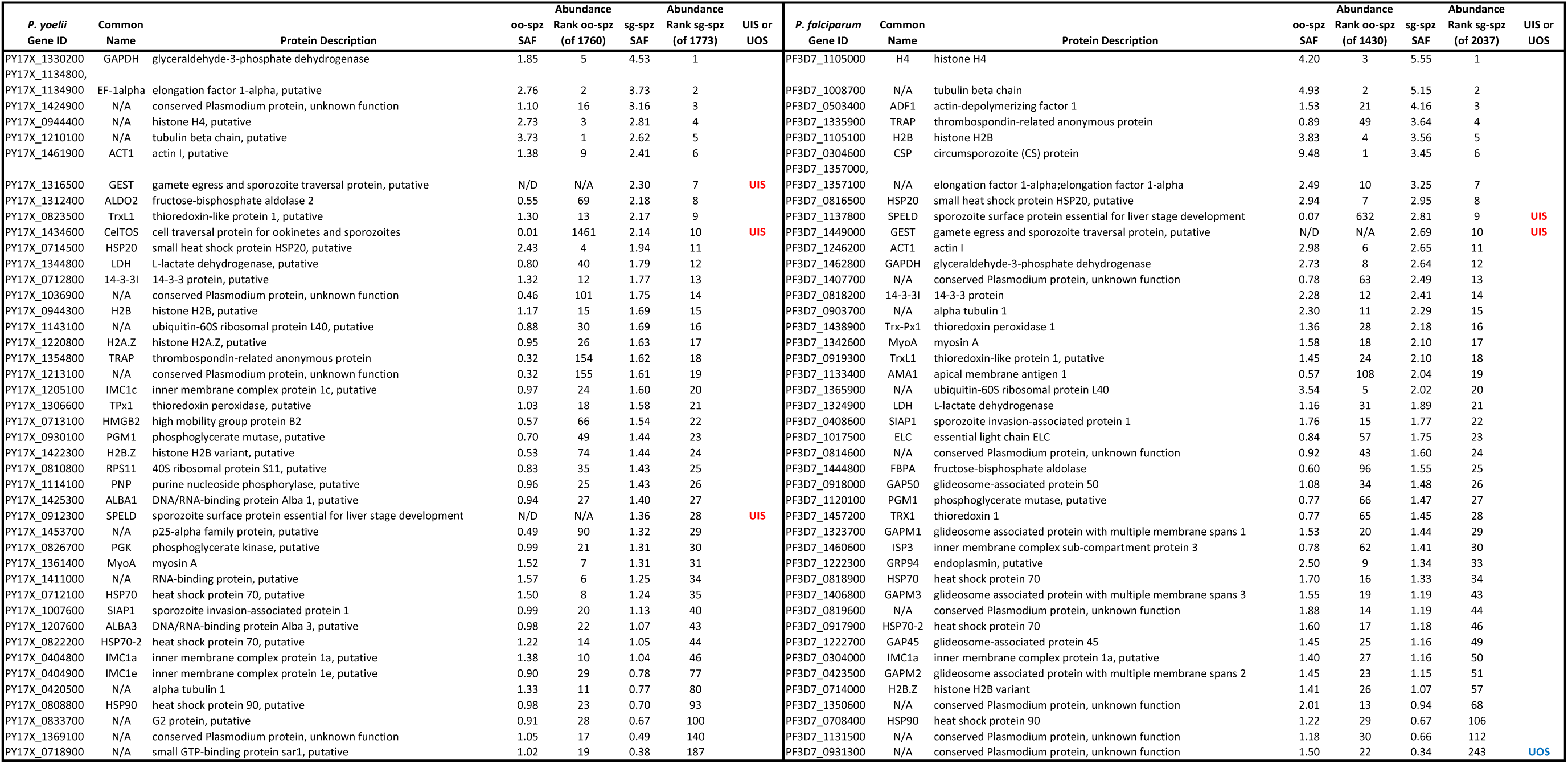
The top 30 most abundant proteins in oocyst sporozoites and salivary gland sporozoites of *P. falciparum* and *P. yoelii*. Proteins are defined as Upregulated in Oocyst Sporozoites (UOS) or Upregulated in Infectious Sporozoites (UIS) proteins as well when their abundance is in the top half of the proteome and their fold change is >6-fold in one sporozoite stage compared to the other. A complete listing of protein abundances using label-free quantification and fold changes between sporozoite stages is provided in Table S1.

With proteomic data from both oocyst sporozoites and salivary gland sporozoites, we have expanded the UIS and UOS designations to proteins that are differentially abundant in one stage or the other. These designated UIS and UOS proteins in *P. yoelii* and *P. falciparum* were identified using the same stringent threshold as was applied to mRNA abundances (>6-fold more abundant in one stage compared to the other). Moreover, this was applied only to the top half of detected proteins of oocyst sporozoites (for UOS proteins) or salivary gland sporozoites (for UIS proteins), as differences in protein abundances quantified by spectral counting methods are most robust among higher-abundance proteins^49^. Based upon this analysis, we identified 30 UOS proteins and 114 UIS proteins in *P. falciparum*, and 65 UOS and 65 UIS proteins in *P. yoelii*. UOS proteins detected in both species include UOS3/TREP, PCRMP2, and PCRMP4 (Figure 3, Table S4), which all have clearly been shown to be expressed in and important to oocyst sporozoites^10, 50, 51^. Similarly, UIS proteins in both species include 6-Cys proteins (P38, P36, P52, B9), CLAMP, GEST, PLP1/PPLP1/SPECT2, PUF2, Sir2A, SPATR, SPELD, TRAMP, and UIS2, which have roles in salivary gland sporozoite infectivity, enable the sporozoite to navigate the host skin and liver, or traverse and productively infect hepatocytes. As with differential expression of RNA, species-specific differences in protein abundance changes were observed, with notable proteins being ApiAP2-SP (*P. falciparum*), and SPECT1, UIS3 and GAMER (*P. yoelii*). These data sets include many of the best-characterized sporozoite proteins, which are also known to be critical to sporozoite maturation and transmission. However, in both species and in both stages, 28-49% of the proteins now defined as UOS and UIS proteins remain uncharacterized and are likely to be important to sporozoite functions in the mosquito vector and the mammalian host.

**Figure 3:**
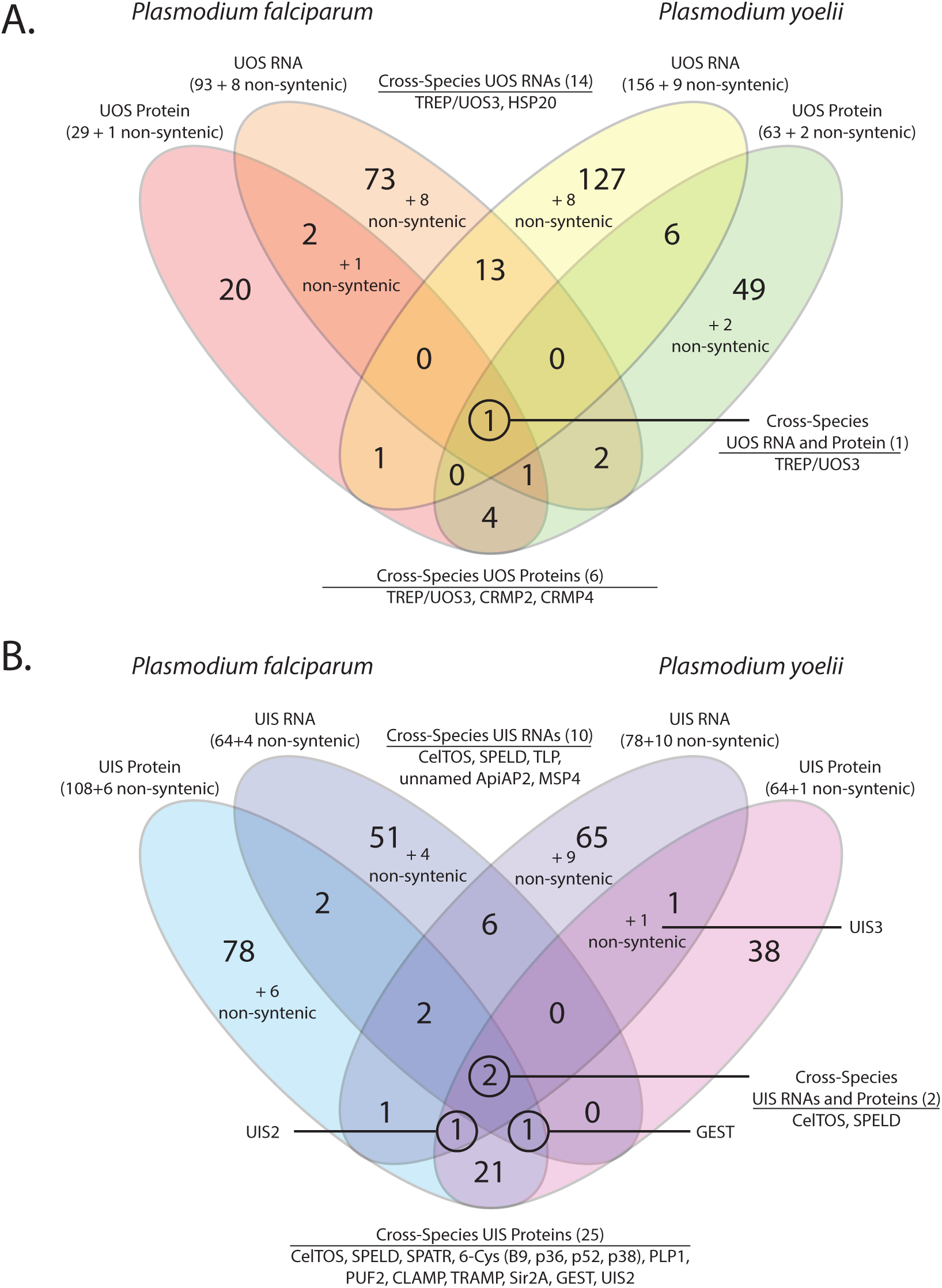
Comparisons of UOS and UIS Gene Products. RNAs or proteins in the top decile of abundance that were also at least 6-fold more abundant in either oocyst sporozoites or salivary gland sporozoites were denoted at UOS or UIS gene products, respectively. Comparisons across molecule types and species for (A) UOS and (B) UIS identify many gene products that are critical/essential to sporozoite development and/or transmission.

### Comparison of UOS/UIS Designations Within and Across Species

When comparisons across species for UOS RNAs and proteins (Figure 3A) or UIS RNAs and proteins (Figure 3B) are made, several key features emerge. First, there are very few gene products that receive the same UOS designations across species (e.g. 14 UOS mRNAs, 6 UOS proteins), few that are both UOS mRNAs and proteins (4 in *P. falciparum*, 7 in *P. yoelii*), and only a single instance of a syntenic gene that encodes a UOS mRNA and protein in both species: TREP/UOS3. However, these proteins are known to be important to the oocyst sporozoite. For example, we identified TREP/UOS3 as well as *Plasmodium* Cysteine Repeat Modular Protein 2 and 4 (PCRMP2, PCRMP4) as UOS proteins in both *P. yoelii* and *P. falciparum*. TREP/UOS3 and PCRMP2 are important for sporozoite targeting to the salivary gland^10, 51^, whereas PCRMP4 is important for oocyst egress^50^. Similarly, relatively few gene products receive the same UIS designations across species (Figure 3B), but those that do include several gene products known to be important to salivary gland sporozoites. For instance, 10 cross species UIS mRNAs and 25 UIS proteins were detected, and include gene products that enable the sporozoite to preserve its infectivity (PUF2), relieve translational repression (UIS2), traverse through host cells (PLP1, CelTOS) and more. Two of these gene products are both UIS mRNAs and UIS proteins in both species: CelTOS and SPELD. While less is known about SPELD, which is found on the sporozoite surface and is essential for liver stage development, CelTOS has been the focus of substantial study and is a promising target for therapeutic interventions^41, 52^. As above, we anticipate that the uncharacterized gene products identified here will play similar roles to those that have already been studied, and deserve prioritization in future work. Together, these data indicate that similar gene regulatory strategies are used by *P. yoelii* and *P. falciparum* sporozoites, but in order to navigate and interact with specific host environments, they must regulate their most abundant gene products differently. Relaxation of the more stringent thresholds used to define UOS and UIS gene products by inclusion of additional expression deciles and/or requiring a lower fold change yields substantially more overlap in the regulated gene products (Table S1).

### Evidence of Two Independent Translational Repression Programs in Plasmodium Sporozoites

*Plasmodium* parasites have adopted the use of translational repression in female gametocytes in a manner analogous to the maternal-to-zygotic transition of metazoans, with translation of stored and protected mRNAs occurring post transmission to the mosquito^18^. However, far less is known about whether a similar, energetically unfavorable regulatory strategy is used in sporozoites. Currently, few transcripts have been shown to be translationally repressed in sporozoites through reverse genetic studies. The best studied example is the *uis4* transcript, which has *cis* control elements located in the coding sequence itself to limit translation of the UIS4 protein prior to transmission^13^. Additionally, a translational repressor, PUF2, has been shown to be essential for the preservation of sporozoite infectivity during an extended residence in the salivary glands^28, 53–55^. Recently, transcriptomic and proteomic data from *P. vivax* sporozoites has indicated that translational repression occurs in this species as well^56^.

In order to identify putatively translationally repressed transcripts in sporozoites, we analyzed our transcriptomic and proteomic data for evidence of highly abundant transcripts for which no protein could be detected. Existing data suggests that translational repression is imperfect, meaning that translationally repressed mRNAs may still produce a detectable amount of protein. Therefore, in these comparisons we used the following highly stringent criteria to define a translationally repressed transcript: 1) transcripts must be in the top decile of mRNA abundance, 2) the corresponding protein must be either undetected or exhibit a disproportionately low abundance (e.g. bottom 50^th^ percentile), and 3) must encode for a protein with detectable tryptic peptides (Table S5). Through comparison of the combined RNA-seq and proteomics datasets, we found that translational repression is extensively imposed upon many of the most abundant mRNAs of both oocyst sporozoite and salivary gland sporozoite stages of both species (Table S6). The extent of translational repression of transcripts in the top decile of abundance is comparable across both species and both sporozoite stages, with each species having transcripts with no evidence (∼40-50% of mRNAs), or no or low amounts (∼68-80% of mRNAs) of protein detected. Specifically, 115 of 164 UOS mRNAs and 71 of 88 UIS mRNAs are translationally repressed in *P. yoelii*, whereas 69 of 101 UOS mRNAs and 50 of 68 UIS mRNAs are translationally repressed in *P. falciparum*. Complete lists of the top 10% most abundant transcripts that are translationally repressed are provided (Table S1).

These datasets also reveal that *Plasmodium* has implemented two discrete and likely orthogonal translational repression programs during sporozoite maturation and transmission. One program imposes translational repression in oocyst sporozoites, which is relieved in salivary gland sporozoites to allow for the production of highly abundant proteins (“TR-oospz to UIS Proteins program”) (Table 3). A second program imposes and retains translational repression upon mRNAs throughout sporozoite maturation (“Pan-Sporozoite TR program”), which may allow for de-repression in the liver stage parasite as in the case of PyUIS4 (Table 4). However, formal demonstration of the full scale of a post-transmission release from the Pan-Sporozoite TR program awaits technical advances to enable total proteomics of early liver stage parasites.

**Table 3:**
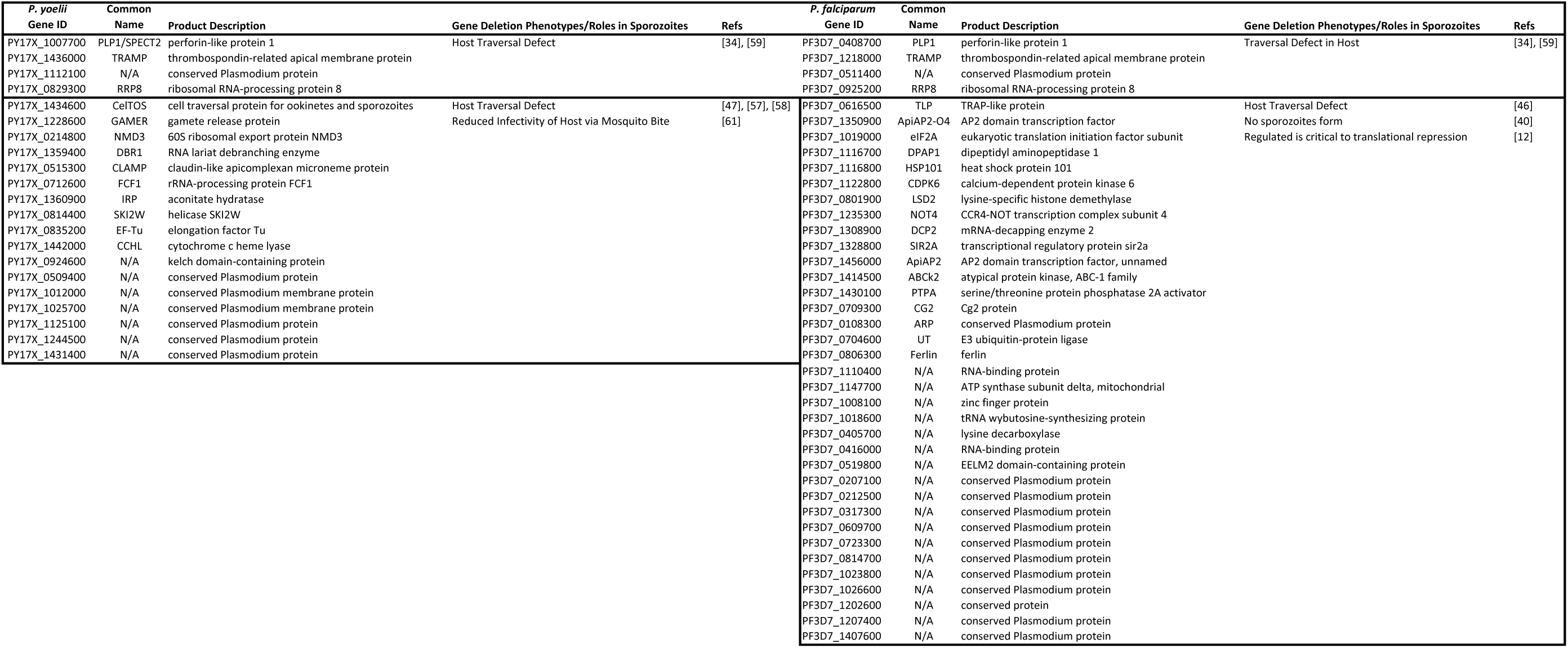
Gene products under the translational repression in oocyst sporozoite to UIS protein regulatory program (“TR oospz to UIS protein program”) in *P. yoelii* and *P. falciparum*. Published gene deletion phenotypes related to sporozoite traversal and infectivity of hosts is noted with references when available. Four gene products (top four rows) are similarly regulated across species.

**Table 4:**
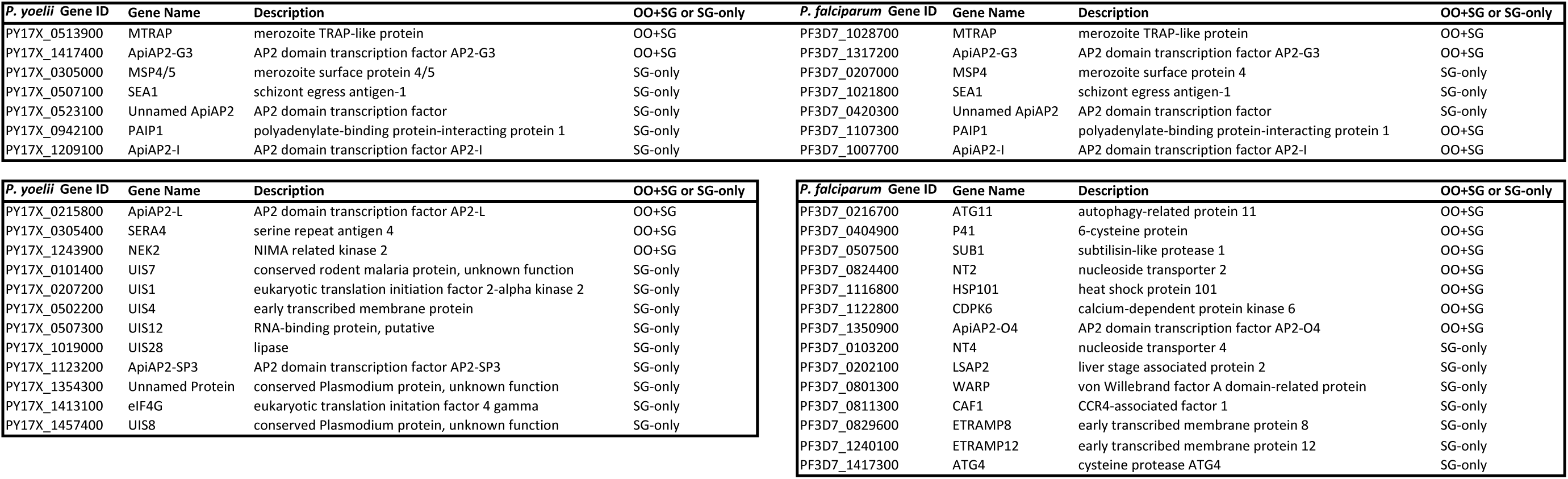
Gene products under the Pan-Sporozoite translational repression program in *P. yoelii* and *P. falciparum*. Gene ID, common gene name, description, and duration of regulation (OO+SG = oocyst sporozoite and salivary gland sporozoite, SG-only = salivary gland sporozoite only). Several gene products (top seven rows) are similarly regulated across species.

Strikingly, in the case of both programs, well-characterized mRNAs are regulated to allow production of their encoded proteins when they are required for the parasite’s activities. For instance, the TR-oospz to UIS Protein program (Table 3) controls production of PLP1/PPLP1/SPECT2, CelTOS, and TLP, which are critical or essential for the sporozoite to navigate the host skin, vasculature, and liver^34, 46, 47, 57–59^. Analyses of the complete TR-oospz to UIS Protein dataset reveal significant GO terms noting roles in the apical invasion complex, the parasite cell surface, movement in host environments, and interaction with and entry into host cells (Table S6). These data are in full agreement with original studies of PLP1/PPLP1/SPECT2 and CelTOS in *P. berghei*, which used IFA, western blotting, and immuno-EM to show that neither protein is present in oocyst sporozoites, but that both become abundant in salivary gland sporozoites^34, 47^. Work on other proteins provide supporting evidence for their expression in and importance to sporozoite functions in the salivary gland and early steps in the infection of the mammalian host^60–62^.

Similarly, the Pan-Sporozoite Translational Repression program affects transcripts that encode for proteins that are known to be important/essential for subsequent stages of the parasite, with notable overlapping regulation of MTRAP, ApiAP2-G3, ApiAP2-I, an unnamed ApiAP2, MSP4, SEA1, and PAIP1 in both species and with similar timing (Table 4). Additionally, in *P. yoelii*, several of the historically defined UIS mRNAs (UIS1, UIS4, UIS7, UIS12, UIS28), ApiAP2-SP3, ApiAP2-L, and others are regulated by this program. In *P. falciparum*, autophagy-related proteins, liver stage associated protein 2, WARP, ApiAP2-O4 and other regulator proteins are affected. This indicates that the sporozoite is capable of immediate regulation of these mRNAs before any significant translation can occur, and is consistent with models that position cytosolic granules near the nuclear pore complex to receive exported mRNAs^63^. Taken together, these data indicate that sporozoites have evolved two overlapping and independent translational repression programs to prepare and remain poised for their next required functions.

### Validation of Translational Repression of Select P. yoelii Transcripts in Sporozoites

To further validate the regulation of select mRNAs by the Pan-Sporozoite TR program, we have used a gold standard, gene-by-gene assessment of wild-type and transgenic *P. yoelii* salivary gland sporozoites by fluorescence microscopy. For this, we have selected genes that exhibit RNA abundances in either the top decile (UIS4 and PY17X_1354300) or the third decile (UIS12), but that by mass spectrometry have protein abundances in the lower half. Previous characterizations of UIS4 in both *P. yoelii* and *P. berghei* have yielded conflicting data on the presence/abundance of this protein using either fluorescence microscopy or mass spectrometry-based proteomics. To address this, we have generated rabbit polyclonal antisera against recombinant PyUIS4 to monitor protein abundance in wild-type sporozoites. By IFA, nearly all salivary gland sporozoites isolated 14 days post-infectious blood meal have no evidence of the UIS4 protein, with few sporozoites having PyUIS4 protein levels that were detectable above background (Figure 4A). This is consistent with a previous report that showed UIS4 protein levels increase over the residence time of *P. berghei* sporozoites in the salivary gland^64^. Together with our current findings, these data are consistent with a robust but incomplete translational repression of UIS4, which becomes increasingly leaky over time in sporozoites, even in the earliest isolatable salivary gland sporozoites. We hypothesize that this might be attributed to our incomplete understanding of how to minimally perturb sporozoites upon extraction from the mosquito.

**Figure 4:**
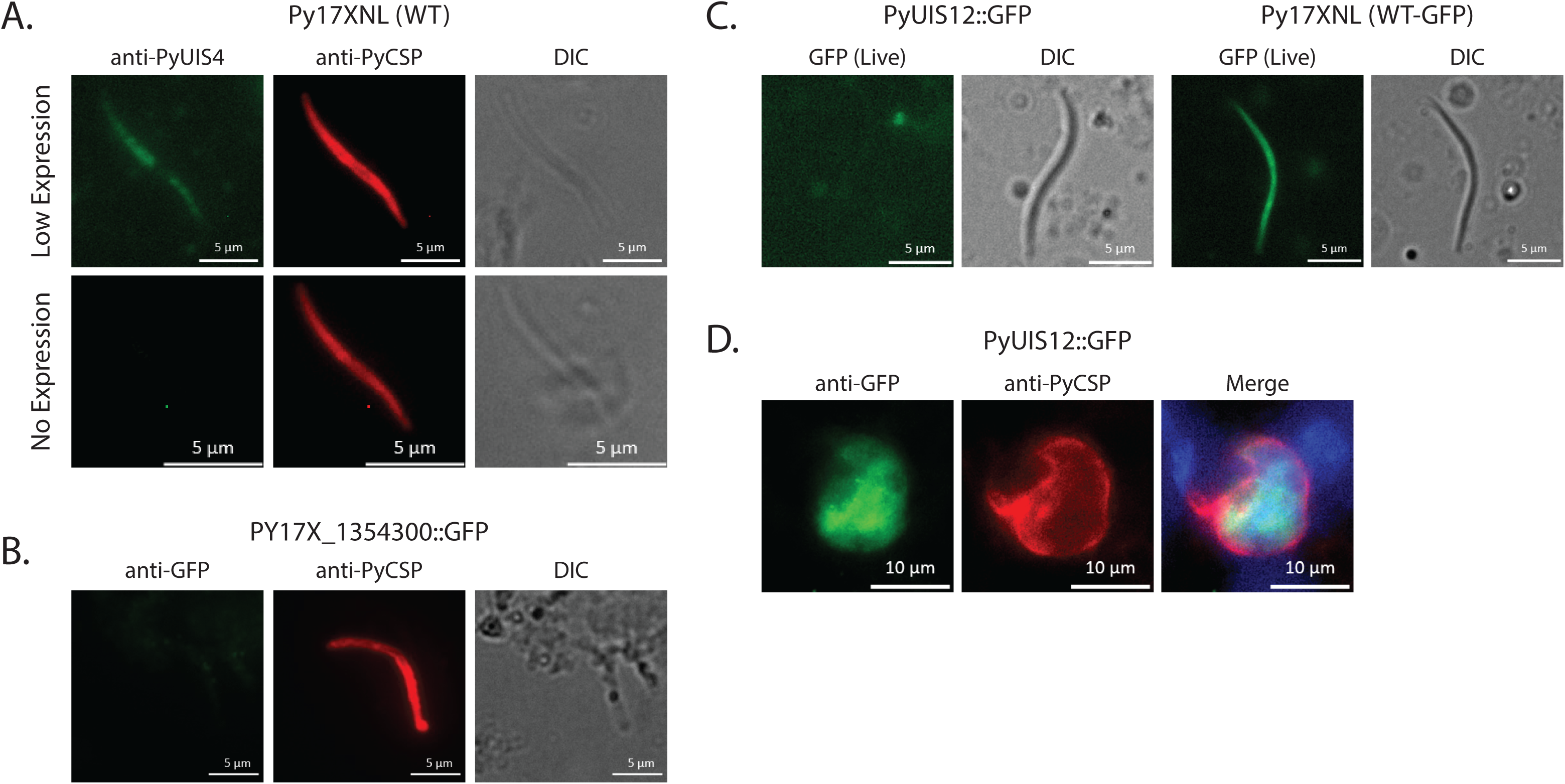
Validation of translationally repressed transcripts in *P. yoelii*. The presence and relative abundance of proteins encoded by translationally repressed mRNAs was assessed by fluorescence microscopy. (A) PyUIS4 expression is largely absent in the population of wild-type sporozoites taken shortly after salivary gland invasion. Very few sporozoites have detectable PyUIS4 protein, which is weak and diffuse. (B) An uncharacterized protein, PY17X_1354300, was modified to encode a C-terminal GFP tag, and protein expression was monitored by IFA. In agreement with the total proteomic data, no protein was detectable. (C) Transgenic parasites expressing unfused GFP (WT-GFP control) or GFP fused to UIS12 were assessed by live fluorescence for evidence of protein expression. No PyUIS12::GFP could be detected above background in sporozoites. (D) However, robust expression of PyUIS12::GFP is detected by IFA post-transmission in 24 hour liver stage parasites.

We have further investigated whether these classifications of translational repression apply to uncharacterized gene products with mRNAs in the top decile of abundance. To this end, we chose PY17X_1354300, which is one of the most abundant mRNAs in *P. yoelii* salivary gland sporozoites (99.5th percentile) but was among the least abundant proteins detected (Table S1). Thus, we created PY17X_1354300::GFP transgenic salivary gland sporozoites, and in agreement with the proteomic data, did not detect the presence of PY17X_1354300::GFP protein using anti-GFP antibodies (Figure 4B). Finally, while we have restricted our definition of translationally repressed transcripts to include only the most abundant mRNAs, it is also likely that less abundant mRNAs are similarly regulated as well. To address this, we have assessed PyUIS12 protein expression in salivary gland sporozoites, as it has high mRNA expression (80th percentile) but was barely detected by mass spectrometry. Using live fluorescence microscopy with PyWT-GFP or PyUIS12::GFP sporozoites, we clearly observed GFP expression in control *P. yoelii* WT-GFP sporozoites, but did not detect UIS12::GFP protein when transcribed from its native locus (Figure 4C). In agreement with translational repression being relieved post-transmission, IFA micrographs clearly show UIS12::GFP expression in the cytosol of 24 hour old liver stage parasites (Figure 4D). Taken together, these data indicate that *Plasmodium* parasites can impose translational repression upon sporozoite transcripts, and can do so beyond what our conservative definition applied to only the top decile of RNA abundance encompasses. However, it is notable that because the consistency and completeness of this regulation varies across individual sporozoites, both global and single cell approaches (such as those applied here) are informative and required to understand this regulatory process.

## Discussion

*Plasmodium* sporozoites offer an intriguing model of parasite infection biology with distinct infectivity profiles in the mosquito vector site of development (oocysts) and site of sequestration for transmission to the mammalian host (salivary glands). Here we report a comprehensive and comparative assessment of the transcriptomes and proteomes of both *P. yoelii* and *P. falciparum* sporozoites. We have captured these gene expression profiles for immature sporozoites from the mosquito midgut (oocyst sporozoites) and mature, infectious sporozoites from the mosquito salivary glands. From these extensive data sets, several important features of transcriptome and proteome regulation can be deciphered that are likely controlling the distinct sporozoite phenotypes in the mosquito vector and mammalian host.

First, these datasets provide a robust classification of transcript regulation across sporozoite maturation at both the mRNA and protein levels. Previous work identified mRNAs upregulated in oocyst sporozoites or salivary gland sporozoites, but relied upon less comprehensive instrumentation and low stringency thresholds. The use of current RNA-seq methodologies and improved genome annotation employed here provides a far more extensive and robust classification of UOS and UIS transcripts, and now does so for both rodent-infectious (*P. yoelii*) and human-infectious (*P. falciparum*) sporozoites (Figure 1, Table 1, Tables S1-3). Moreover, we have also assessed these parasites for large-scale changes in protein abundance through mass spectrometry-based proteomics (Figure 2, Tables 2, S1, and S4). Together, these data strongly align with the expression levels and timing reported for individually studied mRNAs and proteins, and will provide the foundation for a systems analysis of the regulatory networks that govern sporozoite infection biology.

Second, we uncovered evidence that extensive translational repression occurs in both *P. falciparum* and *P. yoelii*, in both oocyst sporozoites and salivary gland sporozoites. In analyzing our data, we first applied rigorous thresholds to interrogate the most abundant transcripts and proteins with the goal of identifying putative targets with the highest possible confidence. We felt that this was prudent, as detection of mRNAs by RNA-seq (which includes sample amplification approaches) is more sensitive than detection of proteins by mass spectrometry (which cannot benefit from sample amplification). Among the top decile of mRNAs by abundance, the encoded proteins for nearly half were not detected at all by mass spectrometry, and the encoded proteins for another quarter were detected at a disproportionately low abundance (Table S6). It is notable that relaxation of these thresholds reveals that translational repression also occurs with less abundant mRNAs, which we also observed through microscopy in the validation of PyUIS12 expression (Figure 4).

Moreover, we find that two translational repression programs appear to be functioning in sporozoites, with some transcripts being translationally repressed in oocyst sporozoites but highly translated in salivary gland sporozoites (“TR-oospz to UIS Protein program”) while others remain translationally repressed throughout sporozoite maturation (“pan-sporozoite TR program”) (Figure 5). In the case of those proteins that have been characterized for their roles in sporozoite maturation and functions in the mosquito and host, clear patterns arise. The TR-oospz to UIS Protein program would provide for the rapid production of proteins in salivary gland sporozoites, and would be well suited for proteins that are needed immediately after transmission for host cell traversal in the skin, vasculature and liver, and/or for productive infection of hepatocytes. In agreement with this, we find several proteins with known roles in cell traversal (PLP1/PPLP1/SPECT2, CelTOS, GAMER, TLP; Table 3). The second Pan-sporozoite TR program, particularly those UIS transcripts that are expressed only in salivary gland sporozoites but that are translationally repressed, would regulate the establishment of a new intrahepatocytic liver stage of infection by allowing for the rapid translation of these mRNAs after hepatocyte invasion. In the absence of robust global proteomic analysis of the early liver stage parasite, which will be exceedingly difficult to achieve, this dataset constitutes a best possible platform from which to assess the liver stage proteome in a candidate-based approach.

**Figure 5:**
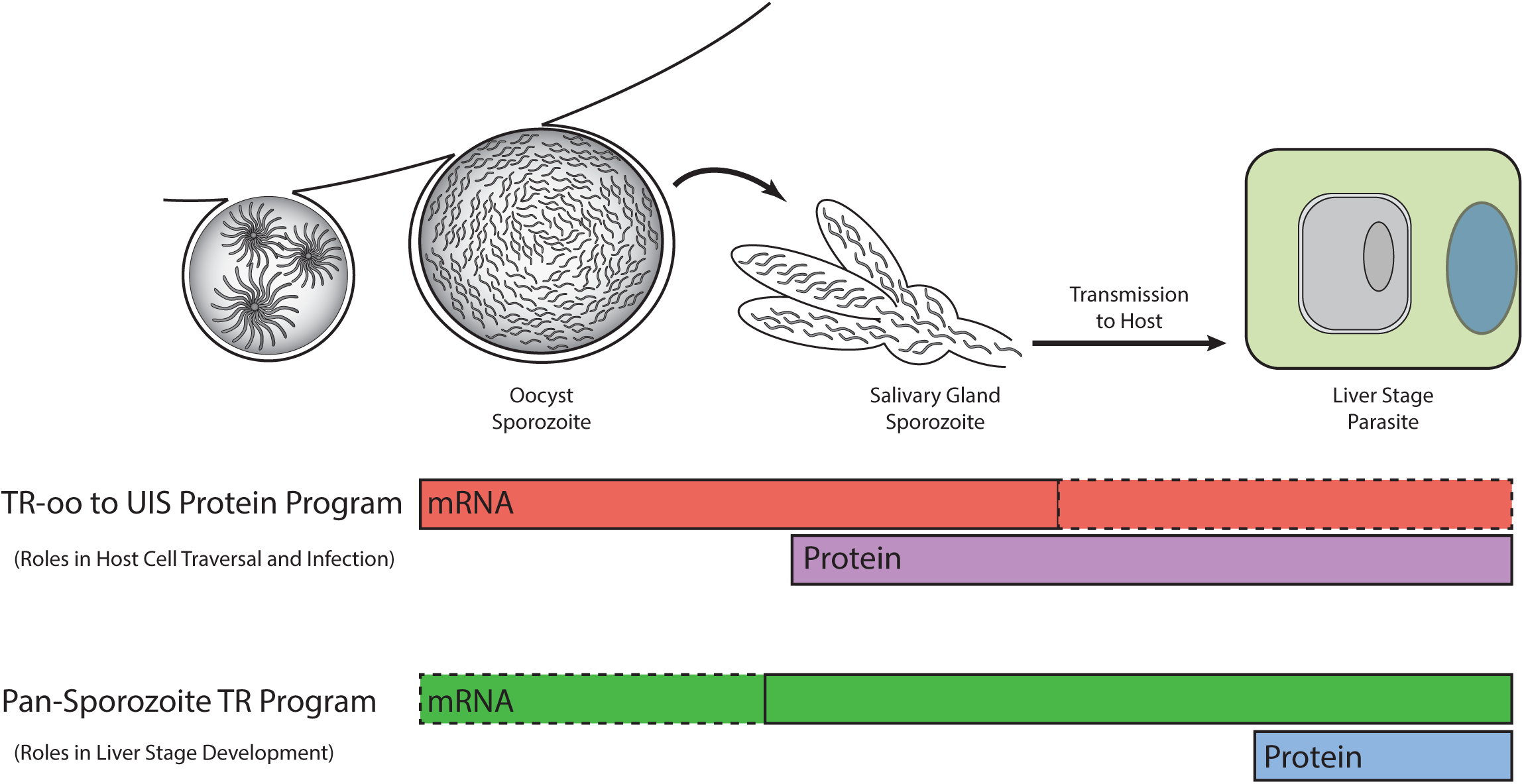
A model for two independent translational repression programs in *Plasmodium* sporozoites. One regulatory program acts upon mRNAs that are highly abundant in oocyst sporozoites to impose translational repression, which is relieved in salivary gland sporozoites and enables protein translation to occur. Transcripts encoding for proteins important to host cell traversal and infection are regulated in this manner. A second regulatory program translationally represses mRNAs throughout sporozoite maturation, which is hypothesized to be relieved following transmission and infection of a hepatocyte to promote liver stage development.

Together, our findings indicate that a multi-tiered, temporal translational repression mechanism is at work in *Plasmodium* sporozoites. This regulatory system aligns with the several windows of functionality that are required for sporozoites’ journey as they egress from oocysts on the mosquito midgut, migrate through the hemocoel, invade the salivary glands, remain there poised for transmission and when transmitted, migrate in the mammalian tissue, traverse cells, and ultimately infect hepatocytes. As the post-transcriptional control of specific mRNAs is energetically unfavorable as compared to *de novo* transcription, we hypothesize that reducing the time between receipt of an external/environmental stimulus and the availability of a protein is critical to the parasite. The use of a TR-oospz to UIS Protein program is straightforward, as it would require a short duration of translational repression until invasion of the salivary glands occurs. However, it is less clear why an energetically unfavorable Pan-Sporozoite program would be activated in oocyst sporozoites, instead of simply transcribing these mRNAs in salivary gland sporozoites. One scenario that could explain the use of both programs is one where the sporozoite invades the salivary gland and then is immediately transmitted, as this would allow immediate responses to both events. While population-level approaches (like those used here) are practical and informative, single-cell approaches should be coupled with them to uncover meaningful differences in the variance of gene expression across individual sporozoites. Enabling technology for single-cell RNA sequencing is currently feasible, and single-cell proteomics is on the horizon.

Finally, new questions emerge from these data. For instance, what are the *trans* factors and *cis* elements responsible for these two likely orthogonal translational repression systems? While several RNA-binding proteins (RBPs) have been implicated in the preparation of salivary gland sporozoites for transmission, specific RBPs have not been associated with specific transcripts in the sporozoite. Moreover, as the TR-oospz to UIS Protein program initiates in the oocyst sporozoite, these experiments must also be pursued in that stage as well. Additionally, at what point are mRNAs governed by the Pan-Sporozoite TR program released for translation? The prevailing model based upon a few gene-specific examples suggests that it should be relieved after hepatocyte infection in early liver stage. However, total proteomic analyses of this stage are not available and will be technically challenging to conduct due to the relatively large proportion of host-to-parasite material. Also, could another program be active in those species that produce latent liver stage forms, called hypnozoite stages? As the commitment to active or latent liver stage forms likely occurs early, having a translational repression program available in sporozoites could allow for this, and could also contribute to determining the ratio of active to latent liver stage parasites that form. Lastly, perhaps the most appealing questions of all revolve around the uncharacterized and under-characterized gene products identified here. These may provide new clues to unappreciated parasite functions, or produce proteins so very different from host proteins that they can be therapeutically targeted.

## Materials and methods

### Production and purification of Plasmodium yoelii and Plasmodium falciparum Sporozoites

Wild-type *Plasmodium yoelii* (17XNL strain) sporozoites were produced as previously described^28^. Briefly, 6-to-8 week old Swiss Webster mice were infected by intraperitoneal (IP) injection of cryopreserved infected blood and were monitored until the peak day of male gametocyte exflagellation. Mice were then anesthetized with an IP injection of ketamine/xylazine and exposed to 150-200 *Anopheles stephensi* mosquitoes for 15 minutes with periodic movements on the cage to promote consistency in the transmission of parasites to the mosquito population. Oocyst sporozoites were collected by microdissection and grinding of mosquito midguts on day 10 post-blood meal, whereas salivary gland sporozoites were similarly collected from salivary glands on day 14 post-blood meal.

All animal care adheres to the Association for Assessment and Accreditation of Laboratory Animal Care (AAALAC) guidelines, and all experiments conformed to approved IACUC protocols at the Center for Infectious Disease Research (formerly Seattle Biomedical Research Institute, Protocol ID #SK-02 to Stefan Kappe) or at Pennsylvania State University (Protocol ID #42678 to Scott Lindner). To this end, all work with vertebrate animals was conducted in strict accordance with the recommendations in the Guide for Care and Use of Laboratory Animals of the National Institutes of Health with approved Office for Laboratory Animal Welfare (OLAW) assurance.

Wild-type *Plasmodium falciparum* (NF54 strain) sporozoites were produced as previously described^31^ by Seattle Children’s (formerly the Center for Infectious Disease Research, Seattle Biomedical Research Institute) and Johns Hopkins University. *P. yoelii* and *P. falciparum* sporozoites were purified by DEAE sepharose and/or two sequential Accudenz gradients as previously described^25, 29, 65^.

### Reverse Genetic Modification of P. yoelii Parasites

*Plasmodium yoelii* (17XNL strain) was genetically modified using conventional, double homologous recombination approaches with the pDEF plasmid vector as previously described^28^. Oligonucleotides used for the creation of targeting sequences are listed in Table S7. The 3’ end of *py17X_1354300* or *pyuis12* (PY17X_0507300) was modified by the addition of the GFPmut2 coding sequence prior to the stop codon. Transgenic parasites were identified by genotyping PCR, with independent transgenic clones being isolated by limiting dilution cloning. Clonal parasites were transmitted to *Anopheles stephensi* mosquitoes to produce salivary gland sporozoites as described above.

### Live Fluorescence and Indirect Immunofluorescence Assays

Wild-type and transgenic *P. yoelii* sporozoites (PY17X_1354300::GFP, PyUIS12::GFP) were subjected to live fluorescence assays and/or an indirect immunofluorescence assay (IFA) (described in detail in^66^) to characterize the extent of translational repression of these candidates. For live fluorescence microscopy of PyUIS12::GFP, freshly produced salivary gland sporozoites placed on glass slides in VectaShield, overlaid with a cover glass slip, and visualized by fluorescent microscopy using a Zeiss Axioscope A1 with 8-bit AxioCam ICc1 camera and Zen imagine software from the manufacturer. Alternatively, fresh salivary gland sporozoites were fixed in 10% v/v formalin for 10 min, and then air dried to a well on a glass slide defined by a hydrophobic coating. Sporozoites were treated for IFA as previously described, using either rabbit polyclonal anti-PyUIS4 (antigen consisting of AA80-224), produced by Pocono Rabbit Farm and Laboratory, Canadensis, PA) or rabbit polyclonal anti-GFPmut2 as primary antibodies and anti-rabbit IgG antibodies conjugated to Alexa Fluor 488 as a secondary antibody.

### Comparative RNA-seq of Oocyst and Salivary Gland Sporozoites

For all oocyst sporozoite and salivary gland sporozoite replicates, RNA was prepared using the Qiagen RNeasy kit with two sequential DNaseI on-column digests, and was quality controlled by analysis on a BioAnalyzer. Barcoded libraries were created using the Illumina TruSeq Stranded mRNA Library Prep Kit, according to the manufacturer’s protocol. Sequencing was conducted on an Illumina HiSeq 2500 using 100 nt single read length on three biological replicates per sample type. The resulting data was mapped to the respective reference genomes (*P. yoelii* 17XNL strain, plasmodb.org v30; *P. falciparum* 3D7 strain, plasmodb.org v30) using Tophat2 in a local Galaxy instance (version 2.1.0). Count files were generated using htseq-count (version 0.6.1) with a minimum alignment quality value set at 30 and a union mode setting. These count files compare the aligned BAM files to a reference GFF file (plasmodb.org v30 for both Py17xNL and Pf3D7) to evaluate the number of reads mapping to each feature, or gene. The count files are combined and compared across conditions using DEseq2 (version 2.11.38), which outputs complete transcript abundance comparisons and performs best among current differential expression tools for three biological replicates. Normalization values for these data are determined by DEseq2 across compared datasets for oocyst sporozoites and salivary gland sporozoites. Statistical metrics utilized by DEseq2 have been previously described^67^. The average number of counts across biological replicates and their standard error of the mean were calculated to allow ranking of transcripts detected over background. Gene ontology terms (components, functions, and processes) were retrieved from PlasmoDB.org (v30). RNA-seq data reported here is available through the GEO depository (Accession #GSE113582).

### Mass Spectrometry-Based Proteomics of Oocyst and Salivary Gland Sporozoites

#### SDS-PAGE fractionation

Purified sporozoites were subjected to SDS-PAGE pre-fractionation and in-gel tryptic digestion essentially as described in^29, 30^. Briefly, samples were electrophoresed through a 4-20 % w/v SDS-polyacrylamide gel (Pierce Precise Tris-HEPES). Gels were stained with Imperial Stain (Thermo Fisher Scientific), de-stained in Milli-Q Water (Millipore), and cut into equal-sized fractions (26 fractions pooled into 13 LC-MS samples for the *P. yoelii* gel and 24 fractions analyzed as 24 LC-MS samples for the *P. falciparum* gel). Gel pieces were then de-stained with 50 mM ammonium bicarbonate (ABC) in 50 % acetonitrile (ACN) and dehydrated with ACN. Disulfide bonds were reduced with 10 mM DTT and cysteines were alkylated with 50 mM iodoacetamide in 100 mM ABC. Gel pieces were washed with ABC in 50 % ACN, dehydrated with ACN, and rehydrated with 6.25 ng/µL sequencing grade trypsin (Promega). After incubating overnight at 37 °C, the supernatant was recovered and peptides were extracted by incubating the gel pieces with 2 % v/v ACN/1 % v/v formic acid, then ACN. The extractions were combined with the digest supernatant, evaporated to dryness in a centrifugal vacuum concentrator, and reconstituted in liquid chromatography (LC) loading buffer consisting of 2 % v/v ACN/0.2 % v/v trifluoroacetic acid (TFA).

#### Liquid chromatography-mass spectrometry

LC and MS parameters were essentially as described previously^29, 31^. Briefly, LC was performed using an Agilent 1100 nano pump with electronically controlled split flow at 300 nL/min (*P. falciparum* sample) or an Eksigent nanoLC at 500 nL/min (*P. yoelii* sample). Peptides were separated on a column with an integrated fritted tip (360 µm outer diameter (O.D.), 75 µm inner diameter (I.D.), 15 µm I.D. tip; New Objective) packed in-house with a 20 cm bed of C18 (Dr. Maisch ReproSil-Pur C18-AQ, 120 Å, 3 µm). Prior to each run, sample was loaded onto a trap column consisting of a fritted capillary (360 µm O.D., 150 µm I.D.) packed with a 1 cm bed of the same stationary phase and washed with loading buffer. The trap was then placed in-line with the separation column for the separation gradient. The LC mobile phases consisted of buffer A (0.1 % v/v formic acid in water) and buffer B (0.1 % v/v formic acid in ACN). The separation gradient was 5 % B to 35 % B over 60 min (*P. falciparum* sample) or 90 min (*P. yoelii* sample). Tandem MS (MS/MS) was performed with a Thermo Fisher Scientific LTQ Velos Pro-Orbitrap Elite (*P. falciparum*) or LTQ Velos-Orbitrap (*P. yoelii*). Data-dependent acquisition was employed to select the top 20 precursors for collision-induced dissociation (CID) and analysis in the ion trap. Dynamic exclusion and precursor charge state selection were employed. Three nanoLC-MS technical replicates were performed for each fraction.

#### Peak list generation

The raw MS data from our previously reported analysis of salivary gland sporozoites^29^ were re-analyzed using the same databases and parameters described here. Mass spectrometer output files were converted to mzML format using msConvert version 3.0.6002^68^ and searched with Comet version 2015.02 rev.0^69^. The precursor mass tolerance was ±20 ppm, and fragment ions bins were set to a tolerance of 1.0005 *m/z* and a monoisotopic mass offset of 0.4 *m/z*. Semi-tryptic peptides and up to 2 missed cleavages were allowed. The search parameters included a static modification of +57.021464 Da at Cys for formation of S-carboxamidomethyl-Cys by iodoacetamide and potential modifications of +15.994915 Da at Met for oxidation, −17.026549 Da at peptide N-terminal Gln for deamidation from formation of pyroGlu, −18.010565 Da at peptide N-terminal Glu for loss of water from formation of pyroGlu, -−17.026549 Da at peptide N-terminal Cys for deamidation from formation of cyclized N-terminal S-carboxamidomethyl-Cys, and +42.010565 for acetylation at the N-terminus of the protein, either at N-terminal Met or the N-terminal residue after cleavage of N-terminal Met. The spectra were searched against a database comprising either *P. falciparum* 3D7^70^ or *P. yoelii yoelii* 17X^71^ (PlasmoDB v.30, www.plasmodb.org^72^) appended with *Anopheles stephensi* Indian AsteI2.3^73^ (VectorBase, www.vectorbase.org^74^), and a modified version of the common Repository of Adventitious Proteins (v.2012.01.01, The Global Proteome Machine, www.thegpm.org/cRAP) with the Sigma Universal Standard Proteins removed and the LC calibration standard peptide [Glu-1] fibrinopeptide B appended. Decoy proteins with the residues between tryptic residues randomly shuffled were generated using a tool included in the TPP and interleaved among the real entries. The MS/MS data were analyzed using the Trans-Proteomic Pipeline (TPP)^75^ version 5.0.0 Typhoon. Peptide spectrum matches (PSMs) were assigned scores in PeptideProphet, peptide-level scores were assigned in iProphet^76^, and protein identifications were inferred with ProteinProphet^77^. In the case that multiple proteins were inferred at equal confidence by a set of peptides, the inference was counted as a single identification and all relevant protein ID’s were listed. Only proteins with ProteinProphet probabilities corresponding to a false discovery rate (FDR) less than 1.0 % (as determined from the ProteinProphet mixture models) were reported.

#### Protein quantification

Relative protein abundance within and between samples was estimated using label-free proteomics methods based on spectral counting. Briefly, the spectral counts for a protein were taken as the total number of high-quality PSMs (identified at an iProphet probability corresponding to an FDR less than 1.0 %) that identified the protein. Spectral counts were quantified using the StPeter program in the TPP^78^. The distributed spectral counts model was used to divide PSMs from degenerate peptides (peptides whose sequences were found in multiple proteins in the database) among proteins containing that peptide in a weighted fashion^79^. Relative protein abundance within samples was ranked using the normalized spectral abundance factor (NSAF)^49, 80^. Relative protein abundance ratios based on spectral counts were normalized and p-values were assigned as previously described^30^. The raw and fully analyzed data files for these mass spectrometry-based proteomic experiments have been deposited in PRIDE (Accession # PXD009726, PXD009727, PXD009728, PXD009729).

#### Prediction of tryptic peptides

The CONSeQuence algorithm was used to identify proteins with detectable fully tryptic peptides with no missed cleavages^81^. A threshold of a Rank score ≥ 0.5 (derived from the combined predictors) was applied, a cutoff that had a sensitivity >70% with a false positive rate <50% when tested on datasets other than the training data as reported by the developers (21813416). Application of this algorithm with this threshold to our published *P. falciparum* salivary gland sporozoite proteome only misidentified 2.9% of all proteins as having no detectable peptides^29^.

## Acknowledgements

We appreciate and acknowledge ongoing scientific discussions with Istvan Albert, Aswathy Sebastian, Manuel Llinás and his research group, as well as advice given on sequencing services provided by the Penn State Genomics Core Facility (University Park, PA). The authors would like to thank the Johns Hopkins Malaria Research Institute Insectary and Parasitology core facilities, particularly, Christopher Kizito for expert rearing of mosquitoes, and Drs. Abhai Tripathi and Godfree Mlambo for production of the *P. falciparum*-infected mosquitos. We are grateful to Bloomberg Philanthropies for support of these core facilities.

Research reported in this publication was supported by Penn State start-up funds (SEL), the National Institutes of Health National Institute of Allergy and Infectious Disease (http://www.niaid.nih.gov/) under award numbers 1K22AI101039 (SEL), 1R01AI123341 (SEL), K25AI119229 (KES), R01AI132359 (PS), R21AI133369 (SHIK), and R01AI134956 (SHIK), by the National Institutes of Health National Institute of General Medical Sciences (www.nigms.nih.gov) under award number R01GM087221 (RLM), by the National Institutes of Health National Center for Research Resources under award number S10RR027584 (RLM), by the National Science Foundation (www.nsf.gov) under award number 0923536 (RLM, KES), and by the Provost’s Postdoctoral Diversity Fellowship from Johns Hopkins University (MS). The content is solely the responsibility of the authors and does not necessarily represent the official views of the National Institutes of Health, the National Science Foundation or the Bill and Melinda Gates Foundation. The funders had no role in study design, data collection and analysis, decision to publish, or preparation of the manuscript.

## Author Contributions

Conducted experiments: SEL, KES, MS, MPW, ENV, KJH, AMM; Analyzed Data: SEL, KES, MPW, PS, SHIK; Wrote manuscript: SEL, KES, PS, SHIK; Funded the work: SEL, KES, PS, RLM, SHIK

## Supporting Table Legends

**Table S1:** Complete transcriptomic and proteomic datasets and their analyses.

**Table S2:** UOS and UIS mRNAs from *P. falciparum* and *P. yoelii*.

**Table S3:** Transcripts in the top seventh to ninth deciles that increase in abundance 10-fold or greater in salivary gland sporozoites as compared to oocyst sporozoites.

**Table S4:** UOS and UIS proteins from *P. falciparum* and *P. yoelii*.

**Table S5:** Tryptic peptide analyses of putatively translationally repressed transcripts using the CONSeQuence algorithm.

**Table S6:** Extended data and GO terms on translationally repressed transcripts.

**Table S7:** Oligonucleotides used in this study.

